# Severe neurometabolic phenotype in *npc1*^-/-^ zebrafish with a C-terminal mutation

**DOI:** 10.1101/2023.02.09.527822

**Authors:** Ana Quelle-Regaldie, Nerea Gandoy-Fieiras, Paula Rodríguez-Villamayor, Sandra Maceiras, Ana Paula Losada, Mónica Folgueira, Pablo Cabezas-Sáinz, Antón Barreiro-Iglesias, María Villar-López, María Isabel Quiroga Berdeal, Laura Sánchez, María-Jesús Sobrido

## Abstract

Niemann Pick disease type C (NPC) is an autosomal recessive neurodegenerative lysosomal disorder characterized by an accumulation of lipids in different organs. Clinical manifestations can start at any age and include hepatosplenomegaly, intellectual impairment, and cerebellar ataxia. *NPC1* is the most common causal gene, with over 460 different mutations with heterogeneous pathological consequences. We generated a zebrafish NPC1 model by CRISPR/Cas9 carrying a homozygous mutation in exon 22, which encodes the end of the cysteine-rich luminal loop of the protein. This is the first zebrafish model with a mutation in this gene region, which is frequently involved in the human disease. We observed a high lethality in *npc1* mutants, with all larvae dying before reaching the adult stage. *Npc1* mutant larvae were smaller than wild type (wt) and their motor function was impaired. We observed vacuolar aggregations positive to cholesterol and sphingomyelin staining in the liver, intestine, renal tubules and cerebral gray matter of mutant larvae. RNAseq comparison between *npc1* mutants and controls showed 249 differentially expressed genes, including genes with functions in neurodevelopment, lipid exchange and metabolism, muscle contraction, cytoskeleton, angiogenesis, and hematopoiesis. Lipidomic analysis revealed significant reduction of cholesteryl esters and increase of sphingomyelin in the mutants. Compared to previously available zebrafish models, our model seems to recapitulate better the early onset forms of the NPC disease. Thus, this new model of NPC will allow future research in the cellular and molecular causes/consequences of the disease and on the search for new treatments.

## 1 Introduction

Niemann Pick disease type C (NPC) is a rare autosomal recessive neurolipidosis with high variable clinical manifestations, from a rapidly fatal neonatal disorder to an adult-onset neurodegenerative disease that may include cerebellar ataxia, supranuclear ophthalmoplegia, cataplexy, epileptic seizures, dystonia, cognitive and psychiatric symptoms (Patterson, 2001; Vanier and Millat, 2003; Vanier, 2010). Liver and spleen involvement is also common. It is estimated that over 95% of the cases are caused by mutations in the gene coding for NPC1, a transmembrane endosomal protein (Vanier, 2010). The remaining cases are due to mutations in the gene coding for NPC2, a small soluble lysosomal protein (Walkley and Suzuki, 2004).

In NPC patients, more than 460 different mutations have been found in different positions throughout the NPC1 gene, which gives rise to a diverse pathogenesis (Polese-Bonatto et al., 2019; Scott and Ioannou, 2004). Near 45% of these mutations are located into a cysteine-rich luminal loop, which is placed between the 855 and the 1098 aminoacids. The abundance of mutations in this region can be cause for decreased stability of this domain, as it may require more disulfide bonds than the other domains for proper folding (Li et al., 2017). Such genetic alteration usually results in protein missfolding and degradation in the endoplasmic reticulum (Scott et al., 2004), and is typically found in patients with severe infantile neurological onset and severe cholesterol-trafficking alterations (Millat et al., 2001; Ribeiro et al., 2001; Park et al., 2003; Gelsthorpe et al., 2008).

*NPC1* codifies a protein of 1278 aminoacids, with 13 transmembrane and three luminal domains: N-terminal, middle luminal and C-terminal (also called cysteine-rich domain). *NPC1* shares homology with regions of the Patched receptor, which binds to the cholesterol-activated protein sonic hedgehog (Davies and Ioannou, 2000; Marigo et al., 1996; Scott et al., 2004). NPC1 and NPC2 participate in transporting recycled lipoprotein-derived cholesterol from late endosomes/lysosomes to the endoplasmic reticulum and plasma membrane (Cologna and Rosenhouse-Dantsker, 2019; Colombo et al., 2021; Li et al., 2016). Impairment of cellular lipid trafficking can lead to accumulation of unesterified cholesterol and other lipids such as sphingomyelin and phospholipids, resulting in endosomal/lysosomal dysfunction (Lloyd-Evans and Platt, 2010; Vance and Karten, 2014). Filipin staining of cultured skin fibroblasts from NPC patients - a test frequently used within the diagnostic protocol - shows accumulation of unesterified cholesterol (Vanier and Latour, 2015). To date there is no curative therapy for NPC, and the only approved treatment for NPC is Miglustat, a drug that inhibits Glucosylceramidase Beta 2 (GBA2) and delays the progression of neurological symptoms (Ridley et al., 2013; Lyseng-Williamson, 2014).

Mouse models of NPC reproduce some of the main pathological disease features, such as hepatosplenomegaly and loss of cerebellar Purkinje neurons and have been used to test drug compounds for therapy (Chen et al., 2020; Maue et al., 2012; Praggastis et al., 2015; Rodriguez-Gil et al., 2020). Due to its biological features and easy genetic manipulation, Zebrafish is commonly used as a model organism in neuronal (Bandmann and Burton, 2010; Quelle-Regaldie et al., 2021a; Quelle-Regaldie et al., 2021b) and metabolic diseases (Hölttä-Vuori et al., 2010; Ka and Jin, 2021). Previous zebrafish models for NPC include morphant knockdown (Schwend et al., 2011; Louwette et al., 2013), and mutant models for *npc1* and *npc2* carrying mutations in the first exons in the N-terminal domain (Lin et al., 2018; Tseng et al., 2018; Tseng et al., 2021; Wiweger et al., 2021). However, zebrafish models with mutations in the C-terminal region of the protein have not yet been developed.

Here, we generated two *npc1* zebrafish lines through CRISPR/Cas9 technology, each with a different mutation at the end of the cysteine-rich luminal loop. To our knowledge, this is the first animal model in which a *npc1* mutation is developed at the beginning of exon 22. We assessed the mutation effects on the fish survival, morphological phenotype, motor performance, pathology, lipid regulation and gene expression.

## 2 Materials and Methods

### Zebrafish care and maintenance

Zebrafish of the AB strain used for the experiments were maintained in the fish facilities of the Department of Zoology, Genetics and Physical Anthropology of the University of Santiago de Compostela. They were maintained at 28ºC with a photoperiod of 14 hours of light and 10 hours of darkness according to previously described protocols (Westerfield, 2000; Aleström et al., 2019). All experiments involving animals followed the guidelines of the European Community and Spanish Government on animal care and experimentation (Directive 2012-63-UE and RD 53/2013) and were approved by the bioethics committee of the University de Santiago de Compostela and the Xunta de Galicia government.

### Mutant generation through CRISPR/Cas9

Zebrafish *npc1* shows 70% sequence identity with its human ortholog. Zebrafish *npc1* (Ensembl ID: ENSDARG00000017180) sequence is composed of 25 exons and codifies for 1276 amino acids. We created the mutation at the beginning of the exon 22 because this is part of the cysteine-rich luminal loop where the most common sites for mutations in *NPC1* related to juvenile and severe onset of the disease are located (Millat et al., 2001; Ribeiro et al., 2001; Park et al., 2003; Gelsthorpe et al., 2008).

Two guide RNAs (gRNAs) placed at exon 22 (which shares 78% sequence identity with the human exon 22) of the *npc1* zebrafish gene were designed with CRISPRscan (https://www.crisprscan.org/):

gRNA1: taatacgactcactataGGAAGACGTAAAACACACTGgttttagagctagaa

gRNA2: taatacgactcactataGGGCTTTGAGCTCTGGTCGGgttttagagctagaa

Each gRNA primer was amplified with the universal primer 5′ - AAAAGCACCGACTCGGTGCCACTTTTTCAAGTTGATAACGGACTAGCCTTATTTTAACTT GCTATTTCTAGCTCTAAAAC - 3′, using iProof™ High-Fidelity DNA Polymerase (Bio-Rad; Hercules, CA, USA) (protocol available upon request). Subsequently, gRNAs were in vitro transcribed with MAXIscript™ T7 Transcription Kit (Thermofisher Scientific; Waltham, MA, USA) and gRNA concentration was measured through NanoDrop® 2000 (Thermo Fisher Scientific) spectrophotometer.

Between 200 and 300 zebrafish embryos at one cell stage were injected with a mix of 15-40 ng/μl of each gRNA and 1.2 μg/μl of TrueCut™ Cas9 Protein v2 (Thermofisher Scientific). Each embryo was injected with 5nl of this mixture. The experiment was repeated three times.

To verify efficiency of the CRISPR/Cas9 protocol, genomic DNA from 10 injected 48 hours post-fertilization (hpf) embryos was extracted with Chelex 100 Resin (Bio-Rad) and PCR amplified with AmpliTaq Gold™ DNA Polymerase (Thermo Fisher Scientific) (protocol available upon request) using the following primers npc1-F: 5′-GGACTTCCCAGTGACATAATGG -3′ and npc1-R: 5′-CCTGGTGCTGATGGAGAAAG -3. The presence of the heteroduplex product of the CRISPR/Cas9 editing was checked in a 5% acrylamide gel.

F0 mutant embryos were raised to adulthood and out-crossed with wt individuals. The resulting F1 generation was also raised to adulthood and checked for CRISPR/Cas9 mutations. Heterozygous *npc1* mutant individuals (around 50% of the fish) were distinguished from wt through genotyping DNA obtained from the caudal fin. Amplified DNA from F1 mutants was used for cloning with the StrataClone PCR Cloning Kit (Agilent; Santa Clara, CA, USA) and then transformed with StrataClone SoloPack (Agilent) in LB-ampicillin plates. Thereafter, 10 colonies of each *npc1* heterozygous mutant were amplified and double checked by 1% agarose gel electrophoresis and Sanger sequencing with 3730xl DNA Analyzer (Thermo Fisher Scientific). Finally, *npc1* heterozygous zebrafish (*npc1*^+/-^) of the F1 carrying the same mutation were in-crossed to obtain F2 homozygous *npc1* (*npc1*^-/-^), which were used for subsequent experiments. All analyses were performed without prior knowledge of the genotype of the individuals, with the exception of RNAseq and lipid analysis in which we needed to know the genotype in order to create the pools of larvae for shipping to the service providers.

### Body length analysis

1 to 3 weeks post-fertilization (wpf) F2 fish from *npc1*^Δ56^ and *npc1*^Δ7^ lines carrying wt, heterozygous (*npc1*^+/-^) and homozygous (*npc1*^-/-^) mutant alleles were used for phenotypic characterization. For motor behavior analysis, body length analysis and histopathological analyses, we have used the same larvae (for histopathological analyses only 2 and 3 wpf larvae) resulting in:

- 1 wpf: from a heterozygous intercross of *npc1*^Δ56^ line, 97 larvae (25 wild type, 47 *npc1*^+/-^ and 25 *npc1*^-/-^); from a heterozygous intercross of *npc1*^Δ7^ line, 66 larvae (18 wild type, 33 *npc1*^+/-^ and 15 *npc1*^-/-^).
- 2 wpf: from a heterozygous intercross of *npc1*^Δ56^ line, 77 larvae (25 wild type, 25 *npc1*^+/-^ and 27 *npc1*^-/-^); from a heterozygous intercross of *npc1*^Δ7^ line, 74 larvae (28 wild type, 28 *npc1*^+/-^ and 18 *npc1*^-/-^).
- 3 wpf; from a heterozygous intercross of *npc1*^Δ56^ line, 86 larvae (28 wild type, 52 *npc1*^+/-^ and 5 *npc1*^-/-^); from a heterozygous intercross of *npc1*^Δ7^ line, 63 larvae (21 wild type, 44 *npc1*^+/-^ and 7 *npc1*^-/-^).

For this purpose, larvae were anesthetized with 0.002% tricaine methanesulfonate (MS-222, Sigma-Aldrich; Saint Louis, MO, USA). Photographs were taken every week with a Nikon Ds-Ri1 camera attached to an inverted fluorescence microscope (AZ100 Multizoom Nikon; Tokyo, Japan). Images were analyzed with ImageJ software (National Institutes of Health; Bethesda, MD, USA) (Abràmoff et al., 2004) and total body length was measured. All fish were genotyped at the end of the experiments.

#### Histopathology

For histopathological analyses, larvae at 2 and 3 wpf from *npc1*^Δ56^ and *npc1*^Δ7^ lines were anesthetized and euthanized (for euthanasia we used an overdose of tricaine methanesulfonate: 0.02%). After taking a tail sample for genotyping, larvae were fixed with 4% paraformaldehyde (PFA) in phosphate buffered saline (PBS; pH 7.4) overnight at room temperature. Zebrafish bodies were dehydrated in ethanol baths of increasing concentrations up to 100%, rinsed in xylene, embedded in individual paraffin blocks, and 3 μm thick sections of the entire larva were stained with hematoxylin-eosin under routine protocol. Larvae sections with representation of all the organs, were chosen for analysis. A light microscopy study was carried with an Olympus BX50 (Olympus life science; Shinjuku, Tokyo, Japan) microscope coupled to an Olympus EP50 digital camera.

### Immunohistochemistry

For immunohistochemical analysis, 3 μm sections of F2 fish of 3 wpf, 5 fish per genotype (wt, *npc1*^+/-^, *npc1*^-/-^) of *npc1*^Δ56^ and *npc1*^Δ7^ lines were dewaxed and rehydrated at room temperature in a humid chamber. Slides were washed with 10 mM phosphate-buffered saline and 0.5% Tween 20 (pH 7.4) in three successive 5 min immersions. Endogenous peroxidase was quenched with Bloxall Blocking Solution (Vector Laboratories, Inc.; Burlingame, CA, USA) and the sections were incubated with a primary rabbit monoclonal recombinant anti-Niemann Pick C1 antibody [EPR5209] (ab134113 Abcam; Cambridge, UK) generated against the C-terminal region of the protein. Immunohistochemical staining was carried out with ImmPRESS HRP Horse Anti-Rabbit IgG Polymer Kit (Vector Laboratories, Inc.), together with the Vector VIP Substrate Kit (Vector Laboratories, Inc.) as chromogen. Negative controls were included in the experiment by using the antibody dilution solution without primary antibody. Sections were counterstained with hematoxylin.

### Survival analysis

For survival analysis, embryos resulting from the cross of heterozygous adults were used. Embryos and larvae were maintained for the desired time, by the end of which surviving fish were euthanized and genotyped. The number of individuals in each group at the start of the experiments were: 1 wpf: 179 individuals: 86 for *npc1*^+/-Δ56^ intercross and 93 for *npc1*^+/-Δ7^ intercross; 2 wpf: 257 individuals: 163 for *npc1*^+/-Δ56^ intercross and 94 for *npc1*^+/-Δ7^ intercross, 3wpf: 188 individuals: 126 for *npc1*^+/-Δ56^ intercross and 62 for *npc1*^+/-Δ7^ intercross ; 4 wpf: 86 individuals: 44 for *npc1*^+/-Δ56^ intercross and 42 for *npc1*^+/-Δ7^ intercross ; and 5 wpf: 40 individuals: 19 for *npc1*^+/-Δ56^ intercross and 21 for *npc1*^+/-Δ7^ intercross . Number of surviving fish for each genotype were annotated and represented using a clustered bar chart of percentages.

### Motor behavior analysis

Motor performance of F2 zebrafish larvae (*npc1*^Δ56^ and *npc1*^Δ7^) at 1, 2 and 3 wpf was quantified with a Zebralab system with a Zebrabox (Viewpoint; Civrieux, France). The quantification software measures the number of pixels moved for each fish in a certain period. For this, 1 wpf larvae were introduced in 96-well plates, while 2 and 3 wpf larvae were placed in 24-well plates. Total larvae movement was measured for one hour, alternating 10 minutes periods of light *vs* dark conditions.

### Statistical analysis of morphological and locomotion data

Statistical analysis and graphs of the morphology and locomotion of the three groups (wt, *npc1*^+/-^ and *npc1*^-/-^) were generated with *GraphPad Prism* version 7 (GraphPad; San Diego, CA, USA), using a non-parametric Kruskal-Wallis test with Dunn’s multiple comparisons test after performing the D’Agostino–Pearson normality test. Statistical significance was established at a p-value < 0.05. Mean with Standard Deviation (SD) are represented in the graphs.

### Lipid staining

Sphingomyelin bodipy (D3522, Thermo Fisher Scientific) and Topfluor Cholesterol (810255P, Avanti; Webster, USA) stock solutions were produced by dissolving 1 mg in 2 ml chloroform, and 20 μl of stock solution was transferred to eppendorf tubes to let evaporate. The powder was then dissolved in DMSO at a concentration of 10 mg/L. In vivo staining of 1 to 2 wpf *npc1*^Δ56^ larvae was carried out for 24 hours at a final concentration of 0.01 mg/L. After staining, larvae were rinsed 3 times with fish water and anesthetized with 0.002% tricaine methanesulfonate.

Confocal photomicrographs were taken with a Leica TCS SPE confocal laser microscope (Leica Microsystems; Wetzlar, Germany). After capturing the images, larvae were used to verify the genotype.

The images were analyzed with ImageJ software (National Institutes of Health; Bethesda, MD, USA). Pixels of the embryo were selected using the tracing tool and the area, the mean and the integrated density of each fish were measured. These measurements were combined in the formula of corrected total cell fluorescence (CTCF): CTCF = integrated density − (area of selected cell × mean fluorescence of background readings). Statistical analysis and graphs were generated with GraphPad Prism version 7, using a Mann-Whitney test after performing the D’Agostino–Pearson normality test. Statistical significance was established at a p-value < 0.05. Mean with SD are represented in the graphs.

### RNAseq analysis

For RNAseq analyses, 6 samples were used (3 wt and 3 *npc1*^Δ56/Δ56^) each of them containing a pool of 15-20 2 wpf F2 larvae, resulting from F1 incrosses. After sampling the end of the tail for genotyping, larvae were immersed in RNAlater (Sigma-Aldrich) and conserved at -20ºC. RNA extraction was performed with miRNeasy Micro Kit (Qiagen; Hilden, Germany). RNA was quantified using NanoDrop® 2000 (Thermo Fisher Scientific), and a total of ∼1000 ng (40 ng/μl) was sent for sequencing. RNA integrity was evaluated in a 2100 Bioanalyzer (Agilent Technologies), and all samples displayed RNA integrity number (RIN) values *>*9 so appropriate for library construction and sequencing. Libraries were sequenced on an Illumina Nova-Seq 150 bp PE run by Novogene (Cambridge, UK). The quality of the sequencing output was assessed using FastQC v.0.11.7 (https://www.bioinformatics.babraham.ac.uk/projects/fastqc/).

Removal of low quality reads, contaminating sequence, low complexity reads (repetitive DNA sequences) and short reads was performed on read pairs using Fastp v.0.20.1 (Chen et al., 2018). Illumina specific adaptors as well as leading and trailing bases with a Phred score < 20 were eliminated; only reads where both pairs were longer than 35 bp post-filtering were retained.

Alignment of filtered reads against the latest version of the zebrafish genome (GRCz11; http://www.ensembl.org/Danio_rerio/Info/Index) and quantification of transcript abundance was performed using kallisto v.0.46.1 (Bray et al. 2016). According to the authors’ description, Kallisto “pseudo aligns” reads to the transcriptome, producing a list of transcripts that are compatible with each read. This software avoids alignment of individual bases against the genome, thus allowing accounting for multi-mapping reads. Aligned reads were assigned to genes based on the latest annotation of the zebrafish genome.

Gene count data were used to calculate gene expression and estimate differential expression (DE) between wt and mutants, using the Bioconductor package DESeq2 v.1.28.1 (Love et al. 2014) in Rstudio v.4.1.2 (R Core Team, 2017). Briefly, size factors were calculated for each sample using the ‘median of ratios’ method and count data were normalized to account for differences in library depth. Next, gene-wise dispersion estimates of gene counts were fitted to the mean intensity using a parametric model and reduced towards the expected dispersion values. Differential gene expression was evaluated using a negative binomial model fitted for each gene, and the significance of the coefficients was assessed using the Wald test. The Benjamini-Hochberg false discovery rate (FDR) correction for multiple tests was applied, and transcripts with FDR < 0.05 were considered differentially expressed genes (DEGs). Hierarchical clustering and principal component analyses (PCA) were performed to assess sample clustering and identify potential outliers over the general gene expression background. Gene Ontology (GO) enrichment analysis of the DEGs was carried out with ShinyGO v0.741 (Ge et al. 2020), using the zebrafish transcriptome as background

### Lipid profiling

Tissue for lipid profiling was obtained from F2 larvae of 2 wpf of *npc1*^Δ56^. All individuals were sectioned into two parts, most of the body and the end of the tail. The body was frozen at -80ºC and the end of the tails were used to determinate the genotype. 5 pools of 20-25 wt and 5 pools of 20-25 *npc1*^Δ56/Δ56^ larvae were employed for the analysis.

The ten samples were sent to Lipotype GmbH (Dresden, Germany) for lipid profiling. Concentrations of phosphatidate, phosphatidylcholine, phosphatidylethanolamine, phosphatidylglycerol, phosphatidylinositol, phosphatidylserine, diacylglycerol, triacylglycerol, sphingomyelin and cholesteryl ester were measured by a mass spectrometer and a final bioinformatic data report was generated by the same company (Surma et al., 2015).

In summary, the amount of a lipid class was calculated by summing the pmol values of the individual lipids belonging to each class. Class amount is then normalized to total lipid content. For normalization, internal standards (one per class) were used. The intensities of the endogenous lipids are normalized to the intensities of the respective standard and based on the used amount of the standard the absolute value of the measured lipid can be calculated. Values are mean of the biological replicates. Statistical significance was calculated by ANOVA after performing the D’Agostino– Pearson normality test. Statistical significance was established at a p-value < 0.05. Mean with SD are represented in the graphs.

## 3 Results

### Generation of mutant *npc1* zebrafish

We verified by acrylamide gel electrophoresis the presence of an heteroduplex product which indicates the efficiency of the CRISPR/Cas9 technique. Two different mutant lines of *npc1* were identified:

- One presented two deletions: an intronic-exonic deletion of 6 bp, starting at amino acid 1081 and a deletion of 50 bp, starting at amino acid 1107 (*npc1*^Δ56^).
- The other presented a 7 bp deletion (*npc1*^Δ7^) starting at amino acid 1121.

Then, the npc1^Δ56^ line was genotyped by agarose gel electrophoresis, while the npc1^Δ7^ line was genotyped by fragment analysis on an automated fluorescent sequencer (3730xl DNA Analyzer). Both mutations were placed at the C-terminal part of *npc1*, at the end of the cysteine rich luminal loop between the transmembrane domains 8 and 9 and were predicted to cause a premature STOP codon (fig. 1).

**Figure 1:**
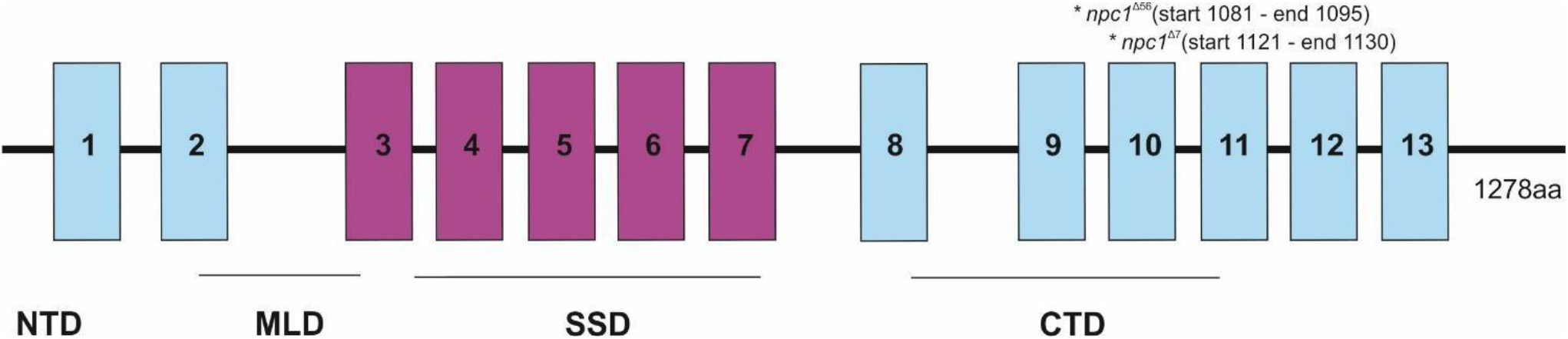
Representation of the *npc1* mutations in the NPC1 protein. 1-13 transmembrane domains were represented and also NTD (N-terminal domain), MLD (Middle luminal domain), SSD (sterol-sensing domain) and CTD (C-terminal domain).

The preliminary characterization of the homozygous mutants (*npc1*^Δ56/Δ56^ and *npc1*^Δ7/Δ7^) revealed similarities in histopathology, survival and locomotion in both lines and therefore these analyses are reported together for both lines by referring to them as *npc1*^-/-^ (see below). The RNAseq and the lipid staining and lipid profiling were only performed in the npc1^Δ56^ line, due to its greater ease of genotyping.

### Morphological and histopathological analysis

*Npc1*^-/-^ and wt individuals showed no apparent macroscopical and histological differences until 2 wpf, albeit only 80% of *npc1*^-/-^ larvae reached this age. The mortality rate was higher at 3 wpf with a survival of 60%. At 4 wpf survival rate of *npc1*^-/-^ larvae was 12% and none of them survived up to 5 wpf (fig.2). From 2 wpf, *npc1*^-/-^ showed a significantly shorter body length (p-value = 0.0343; mean length: wt: 4155 μm ± (SD): 717; *npc1*^+/-^: 3998 μm ± 595.3; *npc1*^-/-^: 3633 μm ± 640.8. From 3 wpf, the smaller size became much more evident (p-value = 0.0037; mean length: wt: 5523 μm ± 889.8; *npc1*^+/-^: 5358 μm ± 1096; *npc1*^-/-^: 4376 μm ± 696.5) (fig.3).

**Figure 2:**
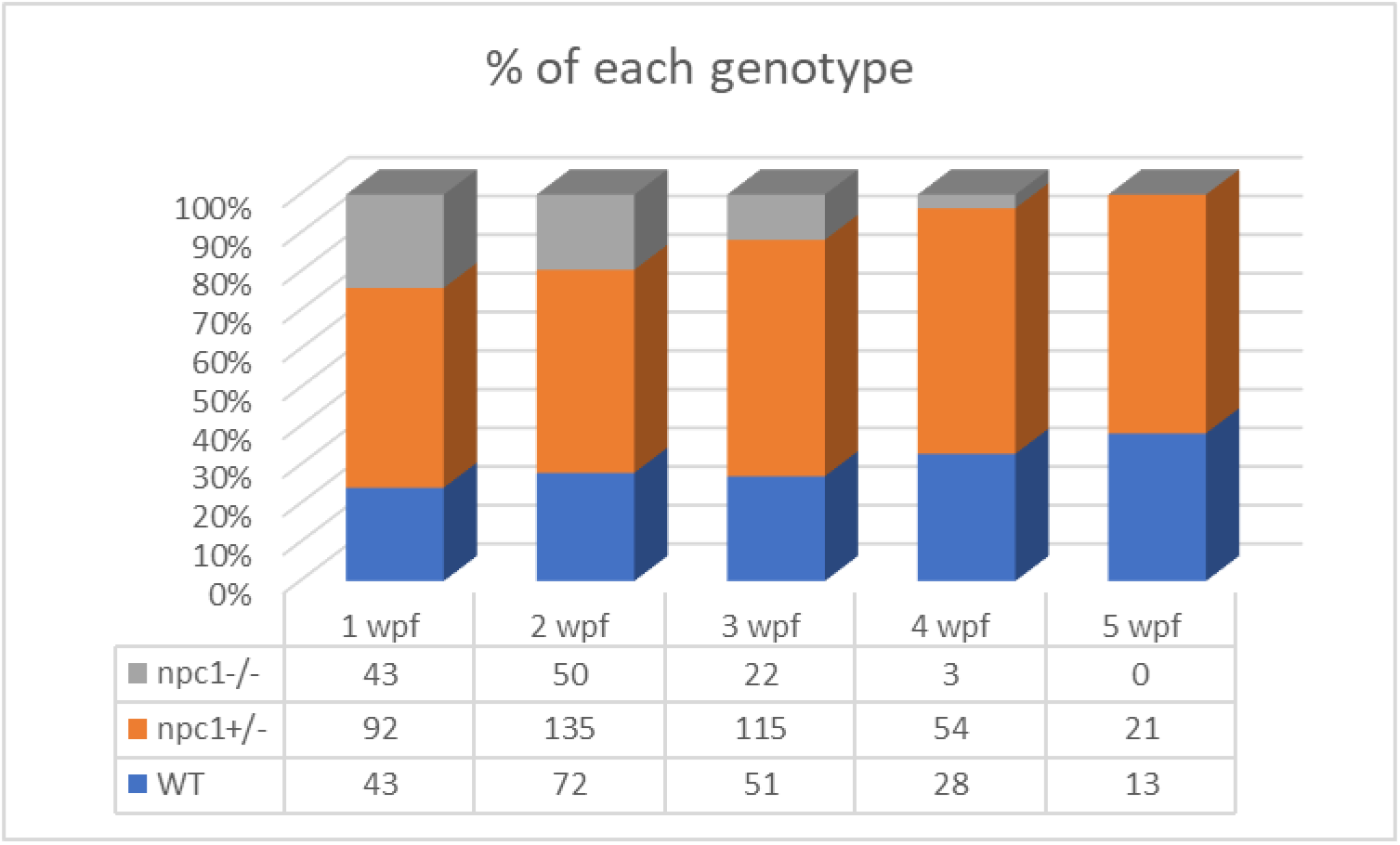
Representation of the percentage of surviving individuals per genotype from 1 to 5 wpf, showing a significant decrease in survival rate of *npc1*^-/-^ larvae from 2 wpf +.

**Figure 3:**
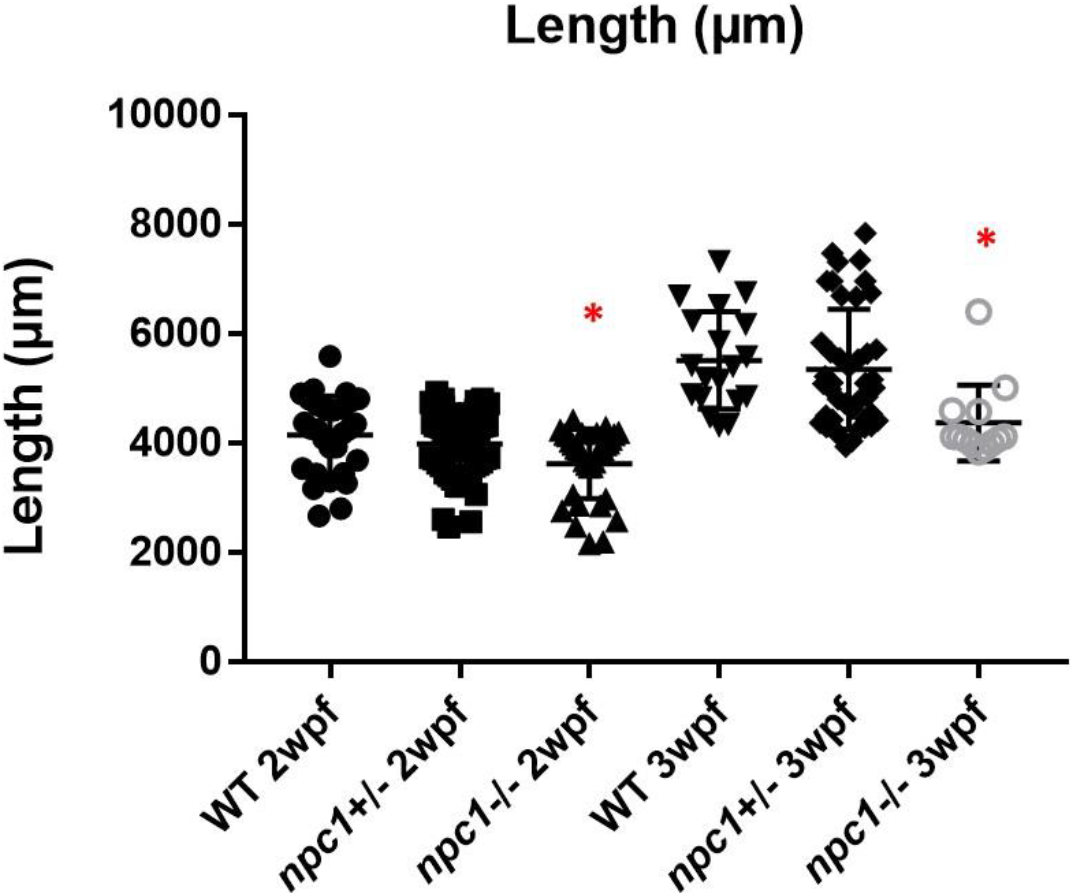
Shorter body length in *npc1*^-/-^ from 2 wpf compared with wt and *npc1*^+/-^ (p-value < 0.05). We compared the fish that had the same age: the 3 groups of 2wpf and the 3 groups of 3 wpf. Statistically significant data in the graphs is indicated with a red *.

A histopathological analysis in 2-3 wpf *npc1*^*-/-*^ larvae revealed well-defined and optically empty intracellular spherical vacuoles, which were compatible with the accumulation of lipids in the liver (fig.4), intestine, renal tubules, and occasionally in the brain (fig.5). Severity of these lesions varied between different *npc1*^*-/-*^ individuals, however the liver was the location of greatest damage in all studied fish. Histopathological grades of vacuolation were assigned as mild, moderate and severe based on an increasing extent of the lesion and the complexity of the changes in the analyzed tissues. The lesions that were found in *npc1*^*-/-*^ larvae were considered moderate to severe degree.

**Figure 4:**
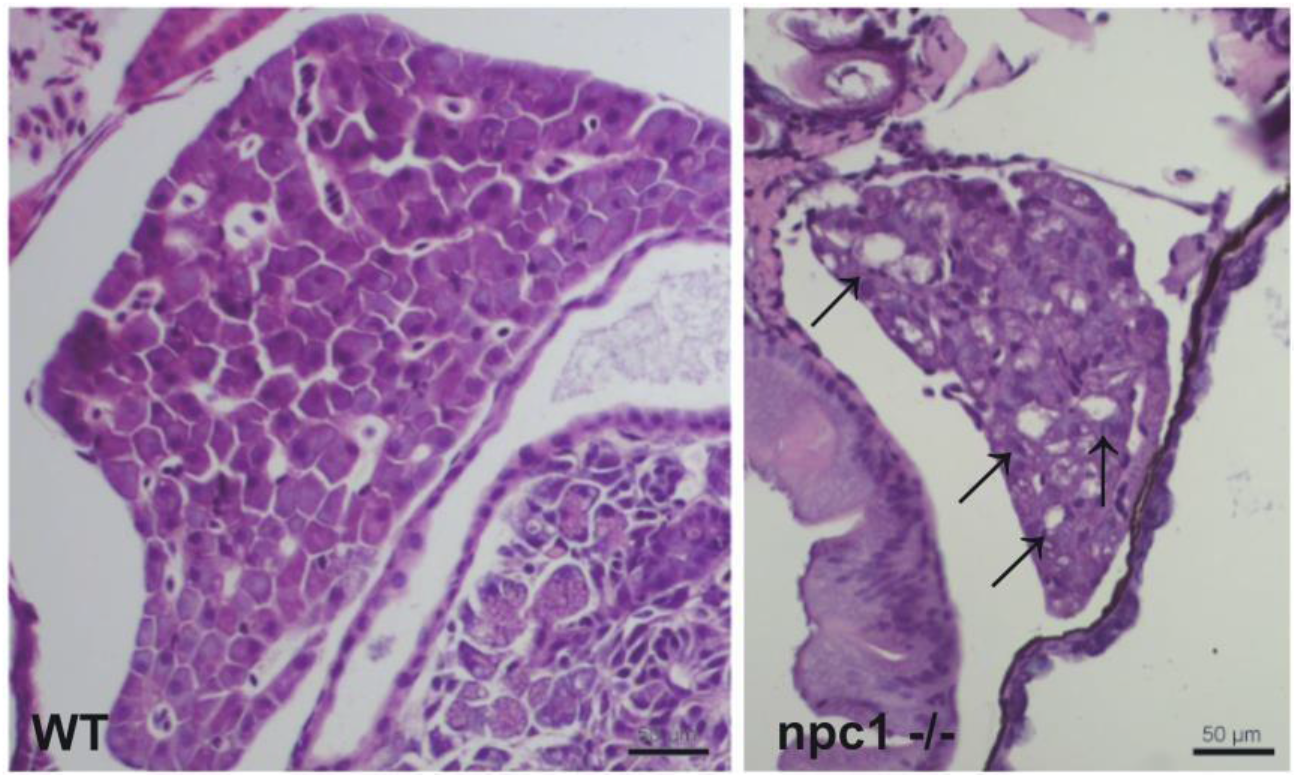
*npc1*^*-/-*^ larvae of 3 wpf have moderate to severe vacuolation and ballooning of hepatocytes (indicated with arrows). Scale bars: 50 μm.

**Figure 5:**
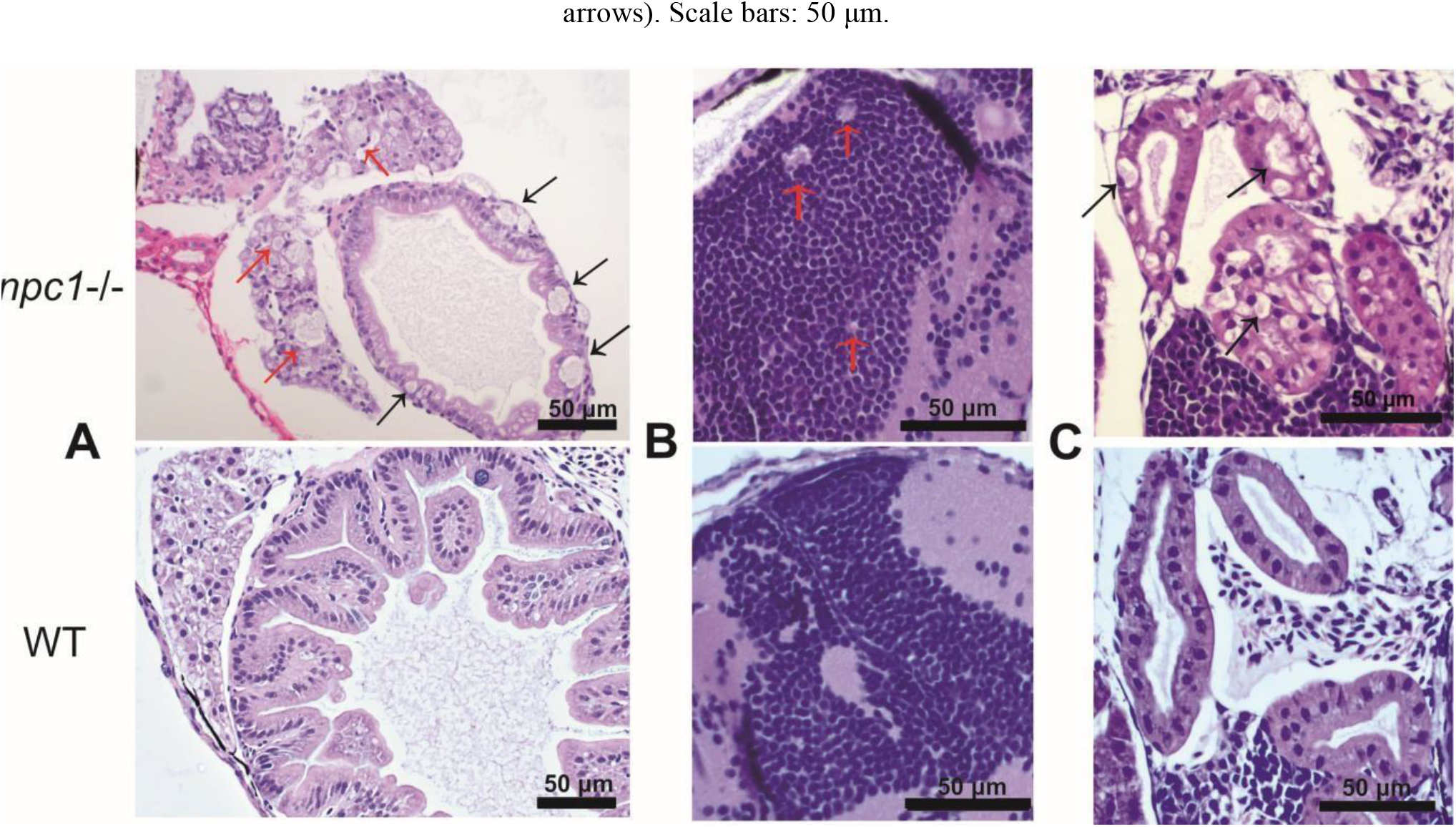
Alterations of different organs in *npc1* ^*-/-*^ and wild type larvae of 3 wpf. A. Intestine and liver. In *npc1* ^-/-^ vacuolation in some enterocytes (intestine: black arrows) and hepatocytes (liver: red arrows) can be observed. B. Gray matter. In *npc1* ^-/-^ vacuolation in the brain gray matter is indicated with red arrows. C. Renal tubules. In npc1 ^-/-^ cytoplasmic vacuolation of renal tubule epithelium is observed, some of the vacuoles are marked with black arrows.

We then aimed to characterize the presence of NPC1 protein in the internal organs by immunohistochemical studies. Our results showed a complete lack of immunostaining in all organs of *npc1* ^*-/-*^ larvae, whereas wt individuals showed immunostaining in the central nervous system, liver, intestine, and renal tubules (fig.6).

**Figure 6:**
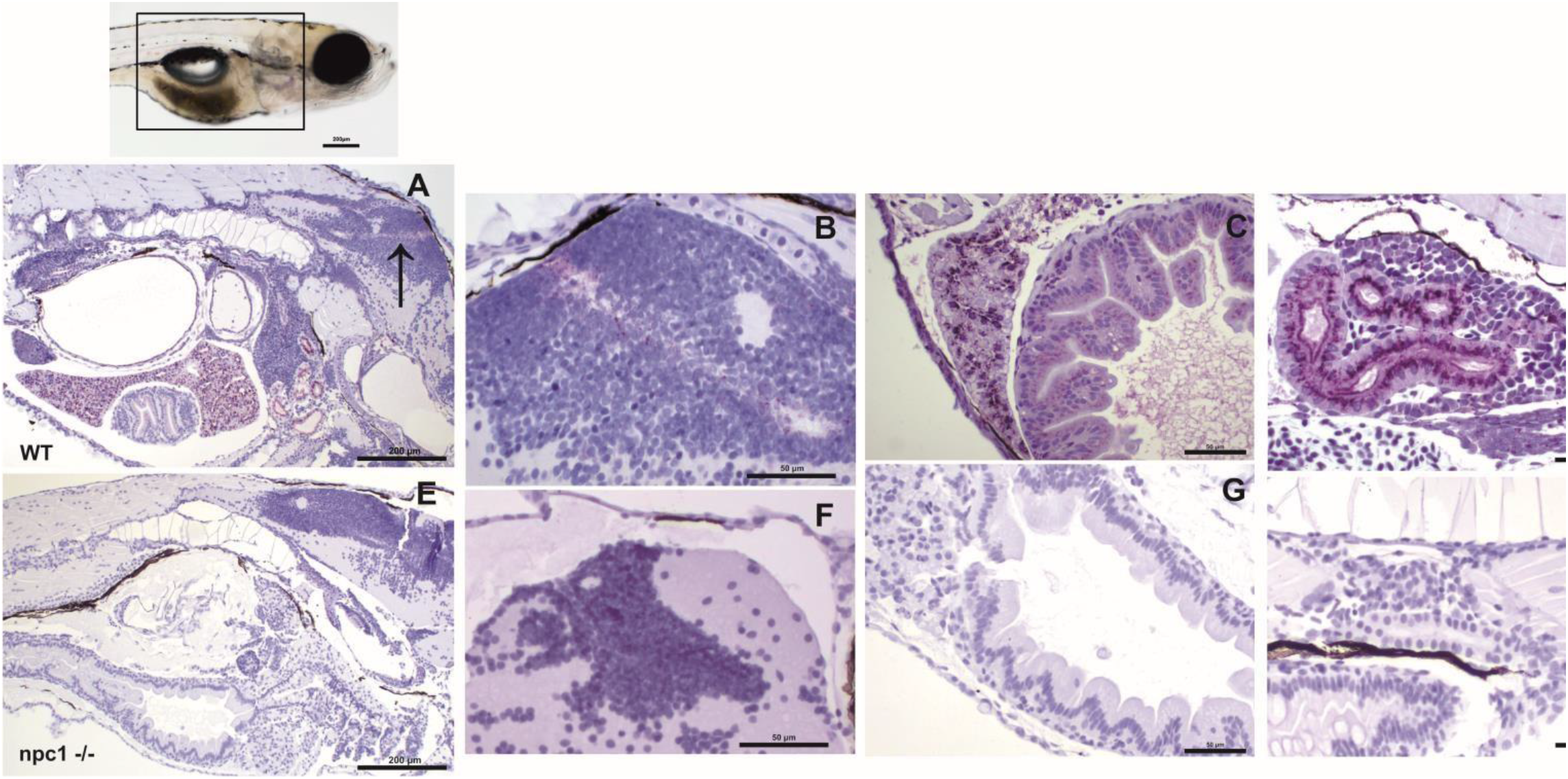
Immunohistochemical study at 3 wpf with anti-NPC1 antibody in wt (A, B, C and D) and *npc1*^-/-^ individuals (E, F, G and H). The square in the top picture indicates the part of the body shown in images A and E. Wt fish (A) were anti-NPC1 positive (indicated by purple staining) in the central nervous system (A) (arrow) and (B), liver (C), intestine (C) and renal tubules (D) whereas *npc1*^-/-^larvae (E) showed no staining in central nervous system (F), liver (G), intestine (G) or renal tubules (H). Scale bars in A and E: 200 μm. Scale bars in B, C, D, F, G and H: 50 μm.

Finally, both cholesterol and sphingomyelin staining resulted in the similar stainings with accumulations of those lipids mainly in the liver of *npc1* ^-/-^ larvae but also in the intestine (fig.7).

**Figure 7:**
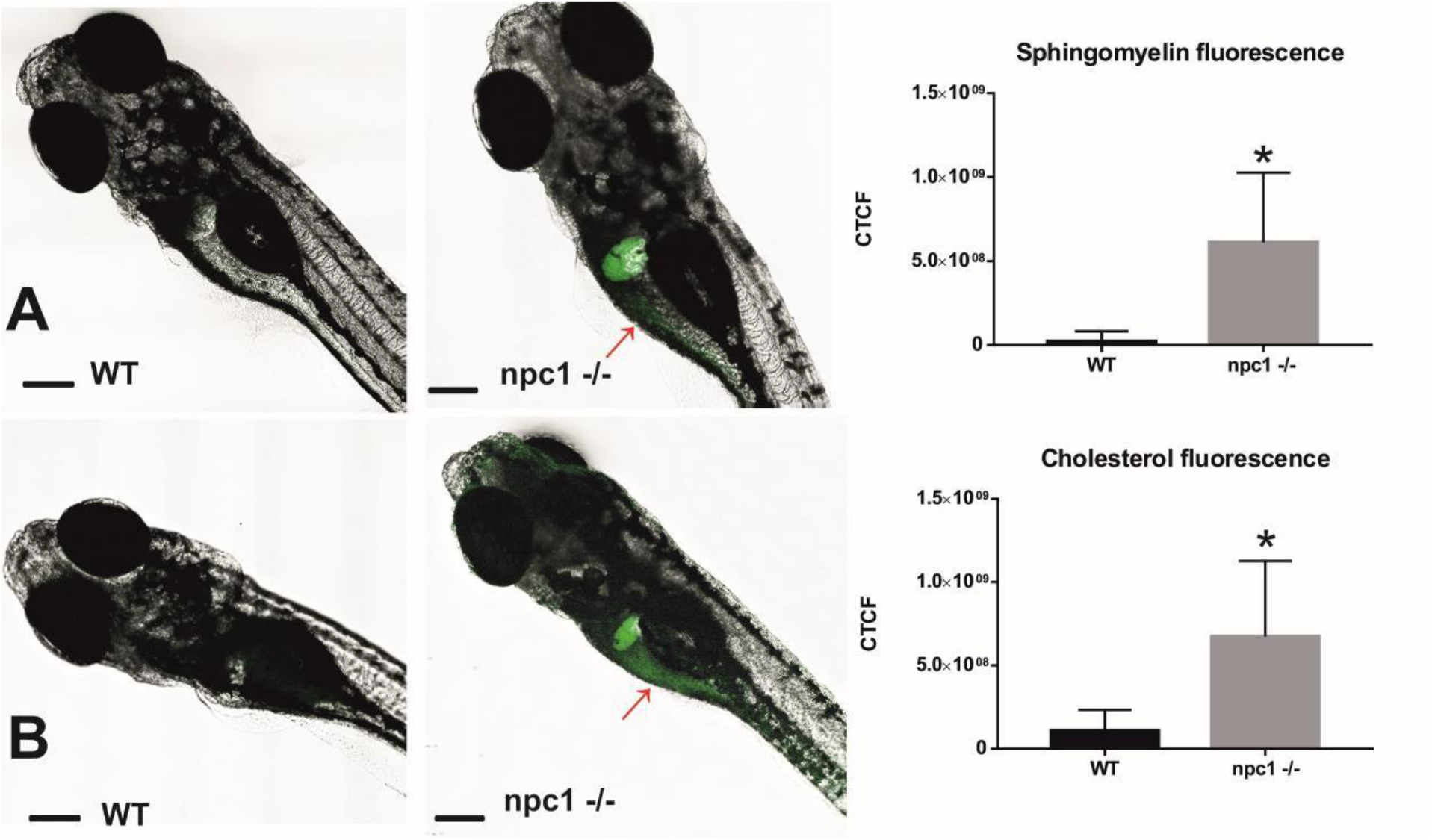
Both sphingomyelin (A) and cholesterol (B) produce aggregations mainly in the liver of the *npc1*^*-/-*^ but also in the intestine (red arrows) at 2 wpf. CTCF test showed increased fluorescence in *npc1*^*-/-*^ animals at 3wpf for both sphingomyelin (A) and cholesterol (B) stainings (p-values < 0.001). Scale bars: 100 μm. Statistically significant data in the graphs is indicated with a *.

### Motor function analysis

At 1 wpf, no differences were found in locomotion between wt, *npc1*^*+/-*^ and *npc1*^*-/-*^ larvae, as measured through comparison of number of pixels moved (not shown). However, at 2 (not shown) and 3 wpf, there was a statistically significant reduction in locomotion of *npc1*^*-/-*^ larvae in comparison with wild type and *npc1*^+/-^ (p-value = 0.0002; mean pixels moved: wt: 133222 ± 142726; *npc1*^+/-^: 150380 ± 173416; *npc1*^-/-^: 68921 ± 64309) (fig.8).

**Figure 8:**
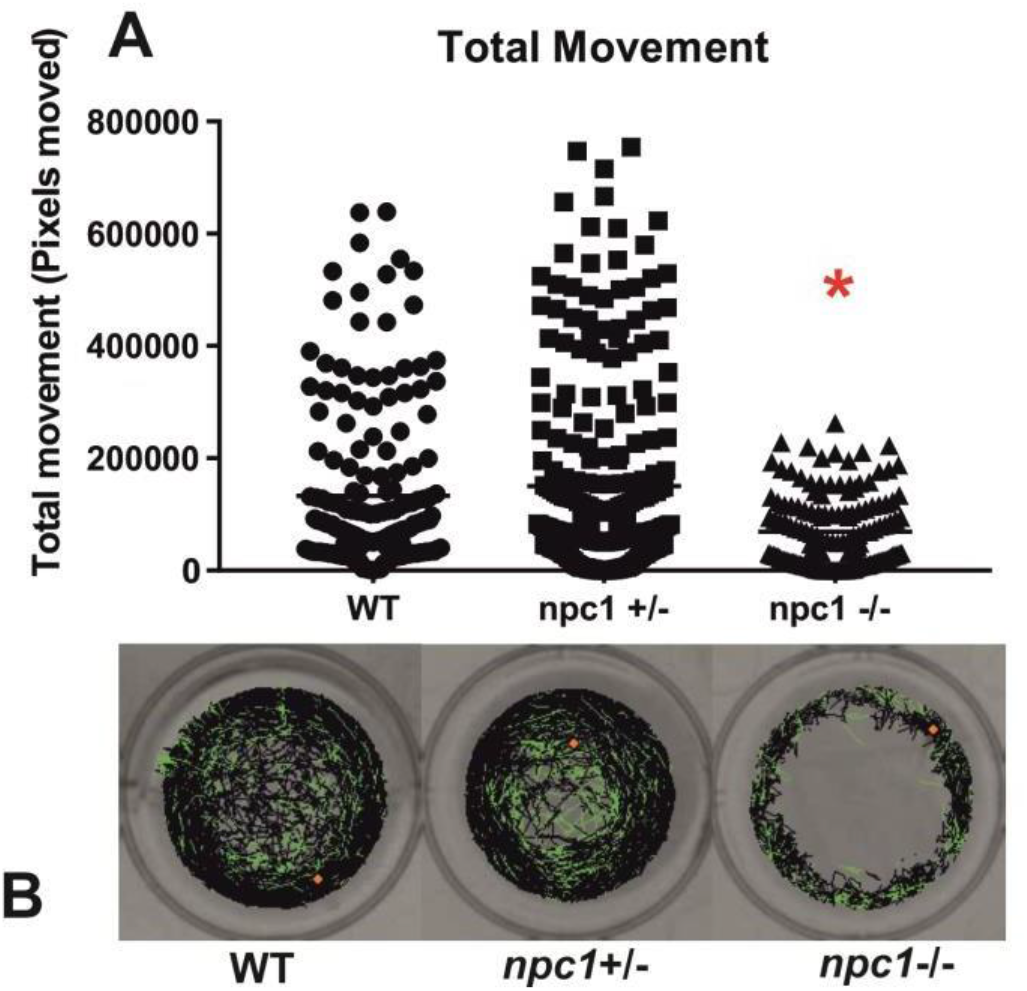
A. *npc1*^*-/-*^ larvae have significant reduction in locomotion compared to *npc1*^*+/-*^ and wt larvae at 3 wpf. p-value = 0.0002. B. Example of locomotion tracks are shown. The orange dot indicated the start position of the larvae. Green color indicated slow movements and black color regular movement. Statistically significant data in the graphs is indicated with a red *.

### RNAseq analysis

RNAseq analysis showed 249 differentially expressed genes (DEGs) between *npc1*^Δ56/Δ56)^ and wt 2 wpf larvae, of which 81 show a higher expression in *npc1*^Δ56/Δ56^ animals and 168 have a reduced expression in *npc1*^Δ56/Δ56^ in comparison with wt (supplemental table 1). Samples of mutant *vs* control individuals did not clearly cluster apart in the principal component analysis (PCA), suggesting the presence of individual variability (fig.9). However, since the *npc1* gene is significantly downregulated in all the *npc1*^Δ56/Δ56^ samples our results can be considered valid to study differential gene expression and perform enrichment analyses.

**Figure 9:**
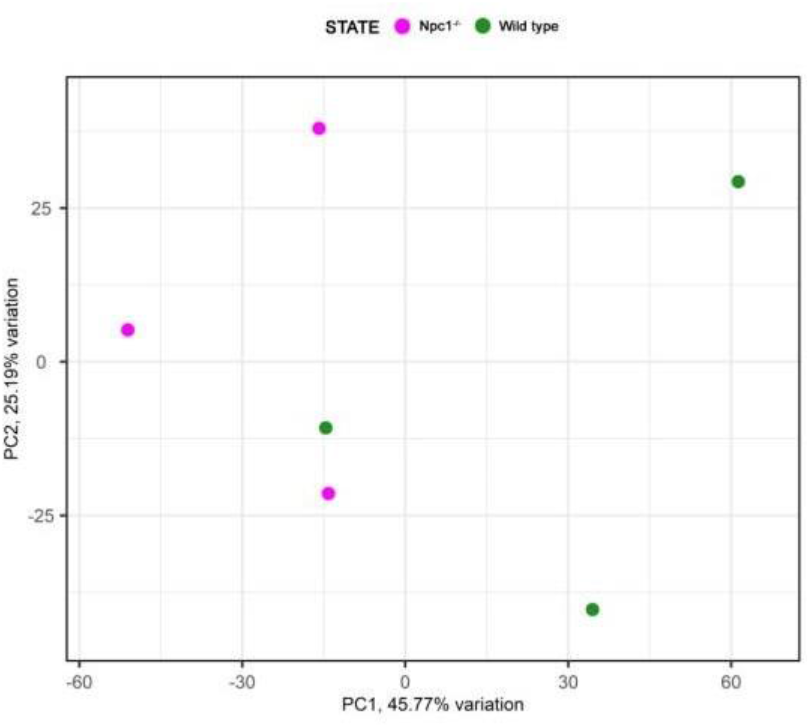
Principal component analysis of the RNAseq samples.

Gene ontology (GO) analysis revealed 357 enriched biological processes, of which 32 involved the *npc1* gene, like ‘lipid homeostasis’ and ‘response to hypoxia’. We also found 111 enriched molecular functions such as ‘transporter activity’, in some of them *npc1* was involved. Other important molecular functions were ‘haptoglobin binding’ and ‘heme binding’ (supplemental table 2).

Out of the 249 DEGs, 14 are involved in transmembrane transporter activity (supplemental table 1). Additionally, we identified 16 DEGs related to lipid transport, modification and metabolic processes; 31 DEGs related to processes of central nervous system such as neural plasticity, axonogenesis, synaptogenesis, myelination and neurotransmitter activity (fig.10); and 12 DEGs related to muscle contraction and development. We also found 12 DEGs related to cytoskeleton activity (supplemental table 1).

**Figure 10:**
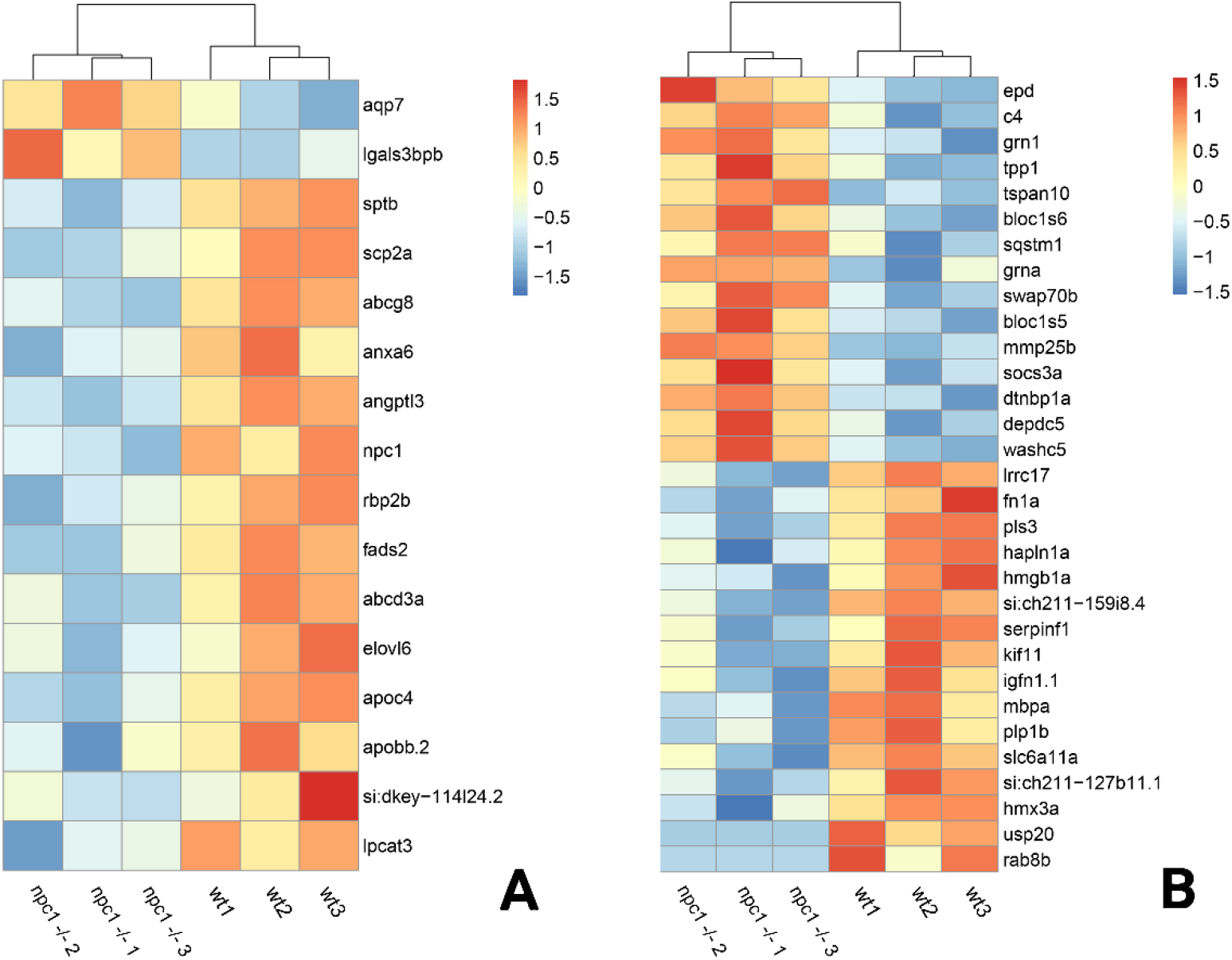
RNAseq analysis revealed differential expression between wt and *npc1*^Δ56/Δ56^ A. Genes involved in lipid synthesis and metabolism. B. Genes related to Central Nervous system processes. Heatmap shows the fold change of selected genes which have significant up- (positive values, orange) or down-regulation (negative values, blue).

Among all DEGs, 20 genes related to ‘collagen extracellular matrix structural constituent’ were down-regulated in *npc1*^Δ56/Δ56^, Furthermore, 17 DEGs related to hematopoiesis and heme oxygenase activity and 9 genes related with angiogenesis were deregulated in *npc1*^Δ56/Δ56^ (supplemental table 1).

### Lipid profiling

5 of the 10 lipid classes of lipids measured showed statistically significant differences in concentrations between *npc1*^Δ56/Δ56^ and wt (fig.11).

**Figure 11:**
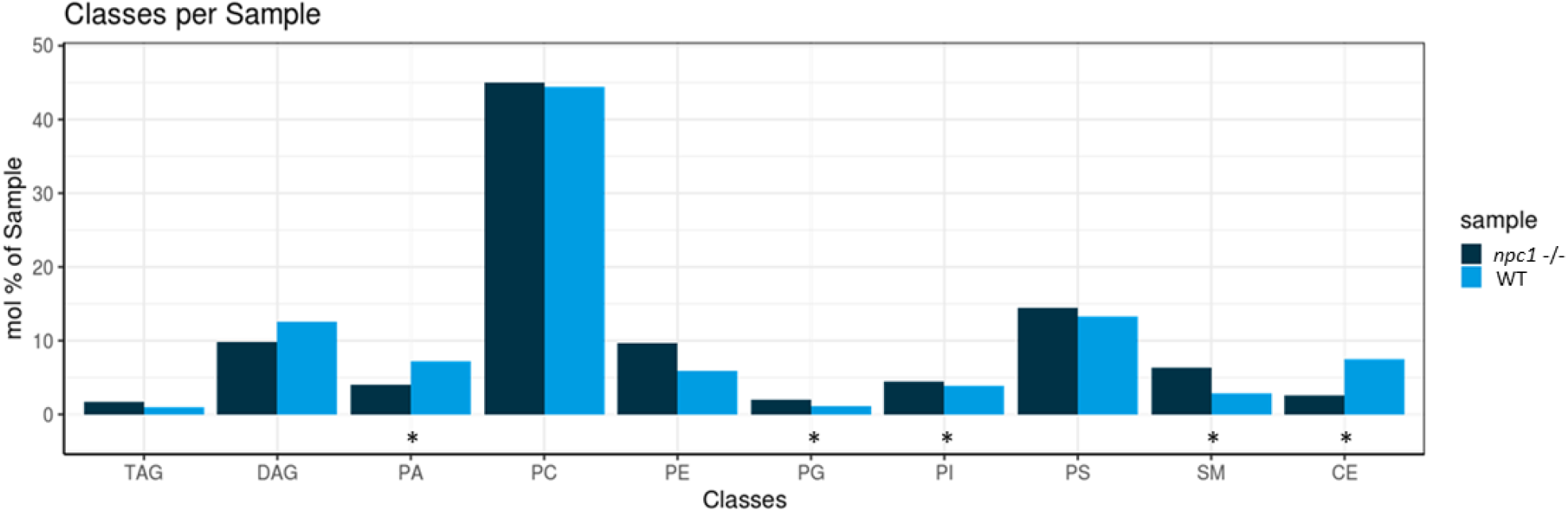
Lipidomic analysis of 10 different lipid classes: TAG (triacylglycerol), DAG (diacylglycerol), PA (phosphatidate), PC (Phosphatidylcholine), PE (phosphatidylethanolamine), PG (phosphatidylglycerol), PI (phosphatidylinositol), PS (phosphatidylserine), SM (sphingomyelin) and CE (cholesteryl esters) between wt (healthy) and *npc1*^-/-^ (disease) revealed statistically significant percentage differences in PA (p-value= 0.036), PG (p-value = 0.0055), PI (p-value =0.038), SM (p-value= 0.002) and CE (p-value = 0.02). Statistically significant data in the graphs is indicated with a *.

Principal component analysis showed a tendency for sample clustering and separation but with variability among samples of the same group (fig.12).

**Figure 12:**
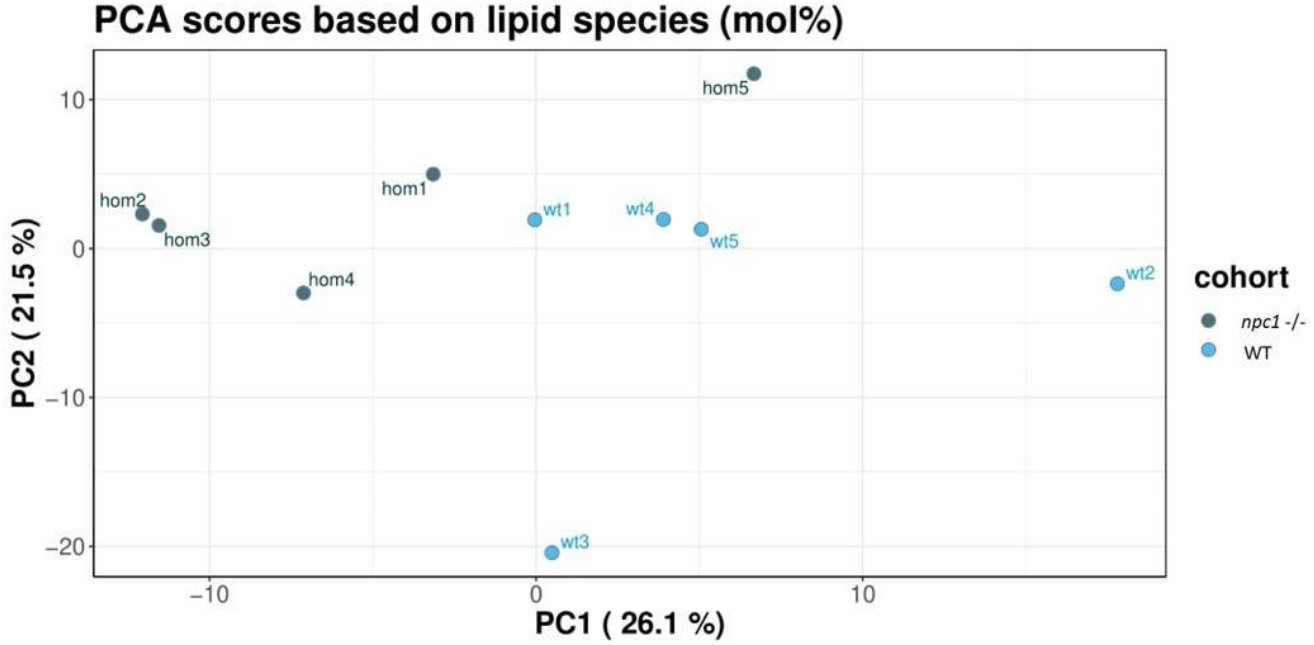
Principal component analysis of the lipidomic samples.

Cholesteryl esters showed an overall statistically significant reduction in *npc1*^Δ56/Δ56^ larvae (p-value = 0.02) compared to wt. 5 cholesteryl species were reduced in *npc1*^Δ56/Δ56^ while another 10 species detected in wt larvae could not be found in *npc1*^Δ56/Δ56^ (fig.12).

Sphingomyelins showed a statistically significant increase in *npc1*^Δ56/Δ56^ samples (p-value= 0.002). 11 sphingomyelin species were found to be more abundant in *npc1*^Δ56/Δ56^larvae than in wt and another 5 species detected in *npc1*^Δ56/Δ56^ could not be found in wt (fig.13).

**Figure 13:**
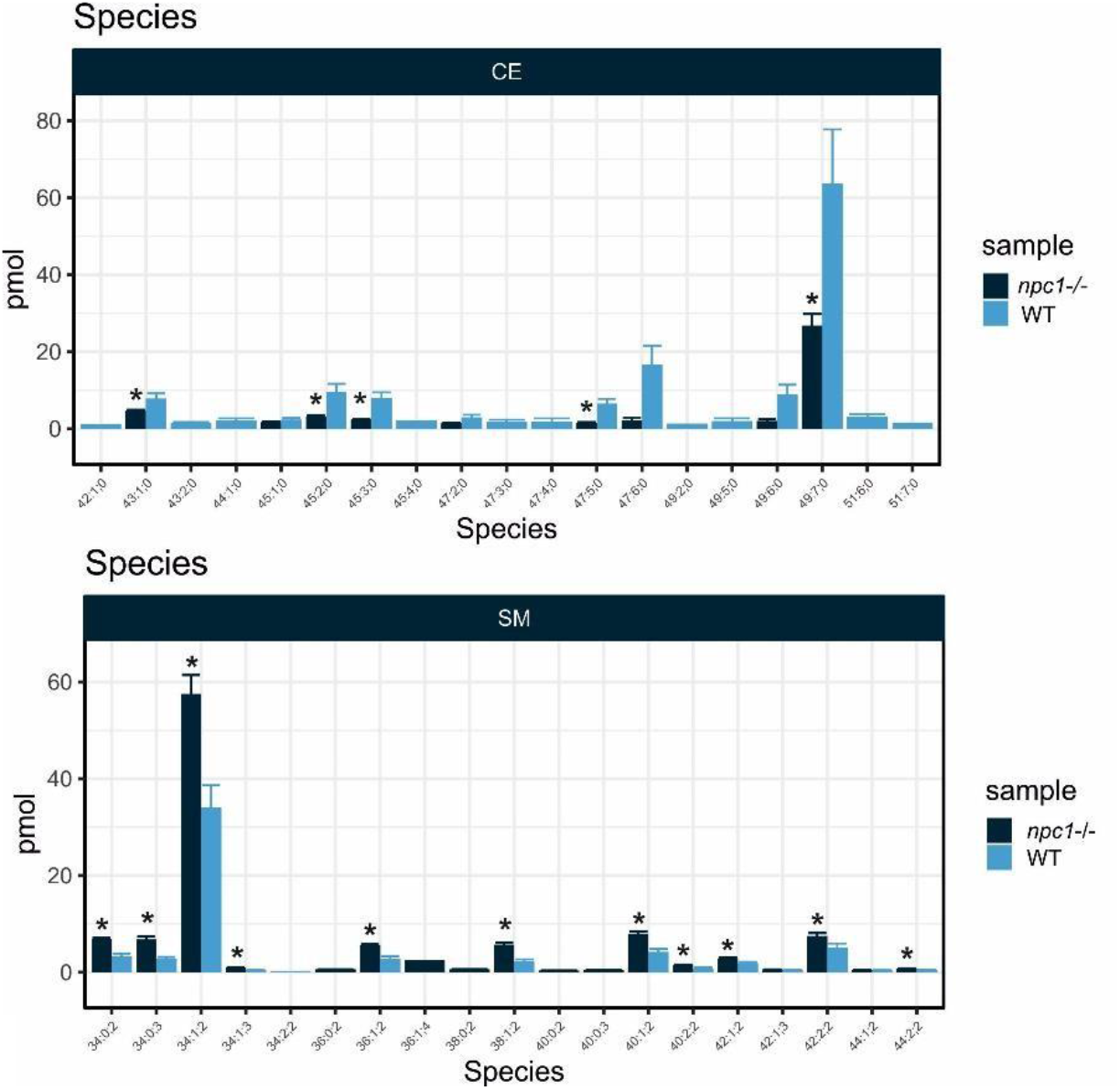
Lipid analysis showed reduction in most species of Cholesteryl Esters (CE) and increase of most species of Sphingomyelin (SM) in npc*1*^Δ56/Δ56^) which is represented as “disease” in the graph in comparison with wt that is represented as “healthy”. Statistically significant data in the graphs is indicated with a *.

Some of the phospholipids analyzed showed statistically significant differences in concentration between wt and *npc1*^*Δ56*/Δ56^ larvae pools. Phosphatidate had a higher abundance in wt than in *npc1*^Δ56/Δ56^(p-value= 0.036). Phosphatidylglycerol and phosphatidylinositol, were reduced in wt compared with *npc1*^Δ56/Δ56^ (p-value = 0.0055 and p-value =0.038, respectively). Concentrations of other lipids (diacylglycerol, triacylglycerol phosphatidylcholine, phosphatidylethanolamine and phosphatidylserine) did not show significant differences between wt and *npc1*^Δ56/Δ56^ larvae.

## 4 Discussion

Animal models are extremely useful for the study of genetic disorders, and essential in the case of rare disorders, such as NPC (between 0.66 and 0.83 births per 100,000 inhabitants) and the high lethality rate (Vanier, 2010).

NPC has been previously studied using animal models, mainly cats and mice but also invertebrates such as the fruit fly and *Caenorhabditis elegans* (Fog and Kirkegaard, 2019). Zebrafish has an *NPC1* gene with a 70% of homology to the orthologous human gene. NPC zebrafish models had been created previously by using morpholinos (Schwend et al., 2011; Louwette et al., 2013) and, more recently, by CRISPR/Cas9 mutagenesis of *npc1* (Lin et al., 2018; Tseng et al., 2018) and *npc2* genes (Tseng et al., 2021; Wiweger et al., 2021). Since there is high variability of mutation type and location leading to human NPC disease, model organisms of this disorder with mutations in different domains are necessary to help elucidate molecular pathogenesis. To our knowledge, this is the first report of an NPC1 zebrafish model with a mutation in the C-terminal domain.

Our NPC models seem to recapitulate the early onset forms of the human disease. Both our *npc1* mutant models carried mutations in exon 22 and died either before or at the beginning of the juvenile stage. This is remarkably different to previous zebrafish NPC mutant models (*npc1* and *npc2* mutant zebrafish had almost the same phenotype), which carried mutations in the first exons and, although they had a reduced lifespan dying between 2-9 months, they survived until at least the start of adult stage (Lin et al., 2018; Tseng et al., 2018; Tseng et al., 2021; Wiweger et al., 2021). Similarly to previous mutant NPC zebrafish models, our mutant NPC zebrafish showed growth retardation resulting in a statistically significant reduced body length. In addition, they showed statistically significant impaired motor function which started at 2 wpf. Locomotion was previous reported to be significant reduced in a *npc2* zebrafish model from 5dpf (Wiweger et al., 2021). Balance defect and trembling was observed but not measured in latter stages of previous *npc1* and *npc2* zebrafish models (Lin et al., 2018; Tseng et al., 2018; Tseng et al., 2021). Early death in *npc1*^Δ56/Δ56^ and *npc1*^Δ7/Δ7^ lines could be mainly due to damage to the digestive organs, but the brain disease that cause decreased locomotion also prevents fish from being able to feed properly.

Additionally, by 2 wpf, we found vacuolated lesions in digestive organs (liver and intestine), as well as in the renal tubules and brain, these being compatible with the accumulation of lipids. Despite similar lesions were also found in previous NPC zebrafish mutants, they were identified at later stages (Lin et al., 2018; Tseng et al., 2018; Wiweger et al., 2021). Therefore, whereas previous NPC zebrafish models appear to undergo a late onset and slowly progressive form of the NPC disease, our *npc1* mutant zebrafish shows a more severe phenotype with an infantile or juvenile onset, resulting in lethality prior to adult stage. Another distinctive feature of our model is the appearance of vacuolar lesions in the renal tubules, which, to our knowledge, had not been previously observed in any other animal model. Curiously, renal failure has been found in a few NPC patients, but all had the adult form of NPC (Philit et al., 2002; Sévin et al., 2007).

*npc1*^Δ56/Δ56^ and *npc1*^Δ7/Δ7^individuals showed a similar pathogenesis to previous *Npc1* mice models that also carry mutations in the cysteine-rich domain of NPC1. These models are characterized by growth retardation, progressive motor impairment, presence of foam cells in the liver, lipid storage, Purkinje cell loss and reduced lifespan (Maue et al., 2012; Praggastis et al., 2015; Rodriguez-Gil et al., 2020). Purkinje cell loss was not studied in our fish but we found CNS alterations (motor impairment, vacuole aggregations in the brain and altered expression of gene pathways related with CNS development by RNAseq). However, these *Npc1* mutant mice had a slowly developing phenotype and had no difference in *Npc1* mRNA levels compared controls and retaining low levels of NPC1 protein. This clearly contrasts with *npc1*^Δ56/Δ56^ zebrafish models, in which we found statistically significant reduction of *npc1* gene expression (RNAseq) and no anti-NPC1 staining across affected tissues (based on immunohistochemistry using antibody against C-terminal) compared to wt larvae. The mice mutations in the cysteine-rich domain have been described as hypomorphic mutations with reduced but not complete loss of gene function. npc1^Δ56/Δ56^ and npc1^Δ7/Δ7^ mutations are also located in the cysteine-rich domain, but they cause a null mutant, which can be explained because NPC1 mutations in cysteine-rich luminal loop were found to cause protein misfolding and degradation in the endoplasmatic reticulum (Scott et al., 2004). This might explain the more severe pathological picture than mice with cysteine-rich domain mutations and other zebrafish models with N-terminal domain mutations (Lin et al., 2018; Tseng et al., 2018; Wiweger et al., 2021).

Previous studies using NPC mutant models have used filipin staining to reveal accumulation of unesterified cholesterol (Lin et al., 2018; Tseng et al., 2018; Tseng et al., 2021; Wiweger et al., 2021). Interestingly, studies about efficiency of filipin test revealed inconclusive results in about 15% of NPC patients (Vanier and Latour, 2015), and recent research has pointed out that filipin test can no longer be considered as the primary tool for NPC diagnosis (Geberhiwot et al., 2018). However, the use of Topfluor Cholesterol and Sphingomyelin bodipy revealed aggregations of both lipids in the liver and digestive organs of *npc1*^Δ56/Δ56^ larvae as soon as 10 dpf.

Lipid profile analysis showed more abundance of sphingomyelin, phosphatidylglycerol and phosphatidylinositol and reduction of cholesteryl esters and phosphatidate in npc1^Δ56/Δ56^ larvae. Sphingomyelin accumulation is typical in NPC human patients. Accumulation of sphingomyelin in NPC cells was shown to inhibit the synthesis of cholesteryl ester and the transport of cholesterol to the endoplasmic reticulum (Wanikawa et al., 2020). Cholesteryl esters function is related to cholesterol transport, and its reduction in *npc1*^Δ56/Δ56^ larvae shows that cholesterol transport is impaired in zebrafish *npc1*^Δ56/Δ56^ larvae. Severe impairment of cholesterol transport was reported before in NPC human patients with cysteine rich luminal loop mutations (Millat et al., 2001; Ribeiro et al., 2001; Park et al., 2003; Gelsthorpe et al., 2008).

Our findings of the lipidomics analysis do not correspond with *npc1* zebrafish mutant created by Lin and colleagues (2018), in which lipid analysis of the liver showed significant differences between wt and *npc1* mutants in profiles of ceramide, diacylglycerol, lysophosphatidic acid, phosphatidic acid, phosphatidylcholine, phosphatidylethanolamine, phosphatidylserine and triglyceride (Lin et al., 2018). We did not measure ceramide, lysophosphatidic acid and phosphatidic acid. However, our measures of diacylglycerol, triacylglycerol, phosphatidylcholine, phosphatidylethanolamine and phosphatidylserine did not reveal significant differences between wt and *npc1* mutants. This could be because our lipid analysis was made with the whole fish instead of liver tissue only, as our fish were young and it would be very difficult to isolate and obtain enough samples of liver tissue for the analysis. In addition, lipid accumulation was shown to be mutation dependent and higher levels of cholesterol and glycolipids were associated with mutations that affected intracellular trafficking (Brogden et al., 2020).

We used RNAseq to carry out whole transcript analysis in *npc1*^Δ56/Δ56^ mutants compared to wt fish. We found statistically significant differences in 249 genes (81 with increased expression and 168 with reduced expression) that were mainly related to transmembrane transport, CNS development, cytoskeleton activity, muscle contraction and development, collagen extracellular matrix structural constituent, hematopoiesis and heme oxygenase activity, angiogenesis and lipid transport and metabolic processes. Differences in genes related with transmembrane transport can be explained as NPC1 was found to function as a transmembrane efflux pump (Davies et al., 2000). NPC neurodegeneration also undergoes cytoskeletal pathology, which has been reported in some murine models of NPC (Bu et al., 2002; Zhang et al., 2004) and cytoskeleton alteration was also previously described in a *npc1* morphant zebrafish model (Schwend et al., 2011). Some previous studies found collagen reduction in some NPC patients as well as in zebrafish and mouse animal models (Louwette et al., 2013; Chen et al., 2020). Moreover, NPC1 has been demonstrated to have a role in hematopoiesis, with some NPC patients showing hematological defects such as thrombocytopenia, anemia and petechial rash and it was also observed in *npc1* zebrafish morphants (Louwette et al., 2013), and angiogenesis has proved to be inhibited by alteration in cholesterol trafficking in NPC individuals (Lyu et al., 2018).

This is the first complete characterization of RNA gene expression in *npc1*^*-/-*^ individuals using the whole body. Previous whole body RNAseq analysis was performed in maternal and zygotic *npc2* mutant zebrafish embryos in which upregulation of lipid transport and metabolism, significant downregulation of genes involved in spinal cord development and genes related with blood vessel endothelial cell differentiation and peripheral nervous system development was observed (Tseng et al., 2021). This was similar to our results because we found altered pathways related to lipid transport and metabolism and blood vessel morphogenesis, although there was a divergence in CNS altered pathways that in *npc1*^Δ56/Δ56^ individuals were found to be axon development and regeneration, neurogenesis and generation of neurons, oligodendrocyte differentiation and neuron projection morphogenesis.

RNAseq analyses at cell and tissue resolution in fibroblasts from NPC human patients revealed altered pathways in cell death and survival, lipid metabolism, small molecule biochemistry, vitamin and mineral metabolism and cellular development, response to incorrect protein, response to endoplasmic reticulum stress, protein folding, protein refolding (Encarnação et al., 2020; Rodriguez-Gil et al., 2021). Most of these pathways with exception of small molecule biochemistry, vitamin metabolism and protein folding and refolding were found to be affected in *npc1*^Δ56/Δ56^ larvae. RNAseq analysis in NPC brain organoids showed that most of the gene differences were those related to nervous system development (Lee et al., 2020)

RNAseq performed in cerebellum and astrocytes of an NPC mouse model (Cougnoux et al., 2021; Han et al., 2021) revealed most discrepancies with *npc1* zebrafish model with altered pathways like immune system process, plasma membrane, protein binding, glutamate transporter, glycosphingolipid biosynthesis, TGF-beta signaling, protein digestion and absorption, cell adhesion molecule, and neuroactive ligand-receptor interaction pathways. Only immune system process, plasma membrane and cell adhesion pathways were altered in *npc1*^Δ56/Δ56^ larvae.

In zebrafish, further studies at tissue level will be necessary to characterize the alteration in the expression of specific genes in the most affected tissues in NPC disease, especially in the brain and liver, and compare them to other models.

## 5 Conclusions

In this study we developed a zebrafish model of *npc*1^-/-^ that will serve to study the effect of a mutation in the last exons of *npc1*, specifically at the beginning of exon 22 which correspond to the end of C-terminal domain, in which the most common mutations of NPC are located. Of note, to date most of the NPC animal models and all zebrafish models were created with mutations in the firsts exons of *NPC1*, less common in the human NPC disease. Due to the large number of mutations found in the NPC1 gene of NPC, it is crucial to develop new models with different mutations, aiming for a better understanding of NPC pathogenesis. Despite further analysis on the CNS consequences of our *npc1*^-/-^ model are needed, we provided extensive evidence of the phenotypic and histological damage, alteration of lipid regulation and gene expression of a suitable NPC model. Previous NPC models showed the zebrafish potential for drug high throughput screening (Tseng et al., 2018). We believe future work with our zebrafish NPC model may help to elucidate new biological markers of the disease and test new therapies to ameliorate NPC symptoms.

## 6 Ethics Statement

The animal study protocol was approved by Ethics Committee of University of Santiago de Compostela (AE-LU-003, ES270280346401).

## 7 Author Contributions

Conceptualization, A.Q.-R.; methodology, A.Q.-R., N.G.-F, P.R.-V., S.M, A.-P.L., M.F., P.C-S. and M.V.-L..; investigation, A.Q.-R., N.G.-F, P.R.-V., S.M, A.-P.L., M.F.; resources, L.S. and M.-J.S.; writing—original draft, A.Q.-R. writing—review and editing, P.R.-V, A.-P.L., M.F., A.B.-I., M.-I.Q.-B. and M.-J.S.; supervision, L.S. and M.-J.S.; project administration, L.S. and M.-J.S.; funding acquisition, M.-J.S. All authors have read and agreed to the published version of the manuscript.

## 10 Supplementary Material

The Supplementary Material for this article can be found online at:

